# Mutators can increase both the rate of occurrence and specificity of nitrofurantoin resistance mutations

**DOI:** 10.1101/2024.10.07.616996

**Authors:** Riannah Kettlewell, Jessica H. Forsyth, Danna R. Gifford

## Abstract

Antimicrobial resistance is a significant global health crisis, with many antibiotics losing effectiveness within a decade of their introduction. The antibiotic nitrofurantoin, however, counters this trend, as it has sustained low resistance rates despite prolonged and widespread use. A key factor behind nitrofurantoin’s success is that resistance requires two independent inactivating mutations in separate genes, *nfsA* and *nfsB*. However, this inherent safeguard may be undermined by elevated mutation rates, a risk that remains unquantified for this antibiotic. Here, we investigated how mutation rates influence nitrofurantoin resistance in *Escherichia coli* using experimental evolution with both low-level and high-level mutator strains, alongside genomic analyses of uropathogenic nitrofurantoin-resistant clinical isolates. Under experimental conditions, increasing mutation rates increased the development of nitrofurantoin resistnace. Moreover, changes in the mutations that occurred also shifted the spectrum of resistance mutations from broad-impact frameshifts and indels to specific amino acid substitutions in the proteins’ active sites. Of the nitrofurantoin-resistant clinical isolates we analysed, nearly 40% harboured disruptive variants in DNA replication fidelity and repair genes*—*a prevalence at the higher end of what is typically observed in uropathogenic *E. coli*. These findings indicate that elevated mutation rates pose a risk to the continued efficacy of nitrofurantoin, highlighting the need for increased monitoring of mutator-driven resistance.

## Introduction

Antibiotic resistance poses profound challenges to healthcare systems; increased health care costs, prolonged illness and increased morbidity and mortality are amongst the many difficulties associated with resistance (Naghavi et al. 2024). Among the many diseases impacted by antibiotic resistance, urinary tract infections (UTIs) are a notable example. Over 150 million UTIs happen each year, with 200,000 resulting in death. Antibiotic resistance is becoming increasingly prevalent in UTIs, with recent studies showing 57%−98% of infections are resistant to at least one antitiotic (Paul 2018; Ahmed et al. 2019; Kot 2019; Ku et al. 2024). The antibiotic nitrofurantoin, however, stands out as an exception among antibiotic treatments. Nitrofurantoin is a WHO Essential Medicine that has been in use as a treatment for UTIs since the 1950’s. Despite this prolonged and steady usage, nitrofurantoin resistance rates have remained consistently low. In 2016, UK guidelines switched from trimethoprim to nitrofurantoin for most UTI cases, resulting in an increase in nitrofurantoin prescribing (Kashouris et al. 2023). Despite this, nitrofurantoin resistance has remained low at about 3%, where in contrast, resistance to the previous front-line drug, trimethoprim, has only slightly decreased from 30% to 26% (Guy et al. 2023; Kettlewell et al. 2024). This has sparked interest in understanding why nitrofurantoin remains effective, and what factors could threaten its continued efficacy (Guy et al. 2023).

Unlike many other antibiotics, nitrofurantoin resistance is almost exclusively aquired through de novo mutation rather than horizontal gene transfer. Moreover, full resistance requires two independent mutations in separate genes (Whiteway et al. 1998; Sandegren et al. 2008). These factors combined likely contribute to the observed low rates of resistance among clinical isolates (Kettlewell et al. 2024). Resistance arises through inactivating mutations in the genes encoding the oxygen-insensitive nitroreductases, *nfsA* and *nfsB* (McCalla et al. 1978). This can occur via various types of mutations, including single-nucleotide variants (SNVs) at key active sites, frameshift-causing indels, disruptions by insertion sequences, and large deletions (Whiteway et al. 1998; Khamari et al. 2022). Crucially, the two genes cannot be inactivated by a single large deletion because they are far apart on the genome (287 kb in *E. coli* K-12, (Vallée et al. 2023); 267-337 kb in other strains) and are separated by hundreds of genes, some of which are essential (16 genes in *E. coli* K-12)(Baba et al. 2006). Mutations in a third gene, *ribE*, are linked to nitrofurantoin resistance, but are likely insufficient alone to confer full resistance (Vervoort et al. 2014).

The reliance on *de novo* mutations is a key factor constraining the prevalence of nitrofurantoin resistance (Kettlewell et al. 2024). However, elevated mutation rates could potentially jeopardize this constraint, increasing the risk of resistance emergence. Under nitrofurantoin treatment, mutation rates may increase due to two specific factors. First, nitrofurantoin is itself somewhat mutagenic, increasing GC>TA transversions and frameshifts (Obaseiki-Ebor and Akerele 1986; Koch and Guillaume 2020). Second, defects in DNA replication fidelity and repair pathways—such as mismatch repair (*mutS, mutH, mutL*), base excision repair (*mutM, mutY*), nucleotide sanitation (*mutT*), and proofreading (*dnaQ*)—can generate ‘mutator’ strains (Fowler and Schaaper 1997; Fukui 2010; Lovett 2011; Fishel 2015; Wozniak and Simmons 2022). Mutators are particularly common among UPEC isolates, occurring in 4–40% of cases (Denamur et al. 2002; Baquero et al. 2004; Couce et al. 2016; Sokurenko 2016). Intriguingly, the mutagenic effect of nitrofurantoin is intensified in some DNA repair mutants, including nucleotide excision repair (*uvrA, uvrB*) and double-strand break repair (*recA, recB*); however, these defects also markedly reduce their MIC to nitrofurantoin (Jenkins and Bennett 1976; Breeze and Obaseiki-Ebor 1983; Ribeiro et al. 2020; Revitt-Mills et al. 2022), complicating the relationship between mutagenicity and resistance.

Although elevated mutation rates have been linked to resistance against various antibiotics (Miller et al. 2004; Jacoby 2005; Sokurenko 2016; Shibai et al. 2023), specific empirical evidence for how elevated mutation rates affect nitrofurantoin resistance is currently lacking. The requirement for two independent mutations to achieve full nitrofurantoin resistance should impede its emergence. However, recent studies indicate that mutators can facilitate multi-locus resistance by enabling the sequential acquisition of multiple mutations (Gifford et al. 2023; Elgrail et al. 2024). Selection for multi-locus resistance can also sometimes drive mutation rate evolution itself (Gifford et al. 2023; Elgrail et al. 2024). Given the widespread use of nitrofurantoin, its mutagenic effects, and the prevalence of mutators among pathogenic *E. coli*, it is essential to investigate the risk mutator strains pose to its continued efficacy and whether nitrofurantoin itself may drive an increase in mutator frequency.

In this study, we examined the role of mutators in nitrofurantoin resistance using two approaches, experimental evolution with laboratory strains exposed to nitrofurantoin, and a genomic analysis of nitrofurantoin-resistant clinical isolates. We first assessed nitrofurantoin resistance evolution in *E. coli* str. K-12 substr. BW25113 with different mutation rates: wild-type (WT), a ‘low-level’ mutator (LM, Δ*mutS*), and a ‘high-level’ mutator (HM, Δ*mutS* + *dnaQ*T15A) (Gifford et al. 2023). Populations with higher mutation rates (LM, HM) evolved nitrofurantoin resistance more readily and had access to ‘better’, more specific nitrofurantoin-resistance mutations achieved via SNVs at active sites in NfsA and NfsB. We did not find a signature of nitrofurantoin mutagenicity in either wild-type or DNA repair deficient strains. We also found no evidence for additional mutation rate evolution among evolved nitrofurantoin resistant isolates. Among clinical isolates, over a third posessed variants in DNA repair genes, including multiple variants with known or suspected mutator phenotypes.

Our results suggest that elevated mutation rates pose a considerable risk for the development of nitrofurantoin resistance. Elevated mutation rates not only increase the likelihood of resistance but also lead to resistance mechanisms that more specifically target nitrofurantoin.

## Materials and methods

### Bacterial strains and media

Bacterial strains used for the selection experiment were wild-type *E. coli* str. K-12 substr. BW25113 (WT), a ‘low’ single-locus mutator (LM, Δ*mutS* (Baba et al. 2006)), and a ‘high’ multi-locus mutator (HM). LM has an approximately 80-fold higher mutation rate than WT. HM arose in the LM genetic background during selection for resistance to a combination of rifampicin and nalidixic acid, during which a spontaneous mutation affecting DNA replication fidelity arose in *dnaQ* (T15A), in addition to mutations in *rpoB* (P564L), *gyrA* (D87G) and other genes (Gifford et al. 2023). DnaQ is the ε subunit of DNA polymerase III, which provides 3′→5′ exonuclease proofreading activity. *dnaQ* T15A occurs in the ExoI motif, one of three regions required for proofreading (Fijalkowska and Schaaper 1996; Lovett 2011; Luan et al. 2013). The mutation rate of HM is approximately 51-fold greater than LM and 1000-fold greater than WT (Gifford et al. 2023).

Cation-Adjusted Muller Hinton Broth (Muller Hinton 2, MH2) was used as the culture medium for all assays (22 g/l; Sigma-Aldrich, UK). For assays requiring solid medium, MH2 was supplemented with 15 g/l agar (BD Biosciences, UK) prior to autoclaving. All incubation was performed at 37°C, the optimal growth temperature for *E. coli*. Nitrofurantoin stock solution was made by dissolving nitrofurantoin powder (TOKU-E, USA) in DMSO (Fisher Scientific, UK) to a final concentration of 8 mg/ml. Stock solutions were filter sterilised using a PTFE 0.22 μm filter (Camlab, UK), aliquotted and stored at −20 °C. Aliquots were thawed prior to adding to growth medium and subsequently discarded. Bacteria were stored in MH2 with 25% glycerol at −80 °C.

### Nitrofurantoin minimum inhibitory concentration assays

Minimum inhibitory concentration (MIC) assays were performed on nitrofurantoin-susceptible WT, LM and HM strains (range tested: 0.25−32 μg/ml), and nitrofurantoin-resistant isolates arising from the selection experiment described below (range tested: 16−512 μg/ml). Glycerol stocks were streaked to singular colonies on MH2 agar plates and left to incubate overnight. MH2 broth was then inoculated with a singular colony for each strain and a 10-fold dilution was performed so that the final concentration of bacterial cells within MH2 broth was 5×10^6^ cells/ml. Cultures were grown in 200 μl volumes within 96-well plates (Nunc, Fisher Scientific, UK). After overnight growth, optical density (OD) readings at 600 nm were taken using a BMG FLUOStar Omega reader (BMG Labtech, Ortenberg, Germany). MIC was recorded as the minimum concentration needed to inhibit growth.

### Selection experiment with nitrofurantoin

To investigate the role of mutation rate in antibiotic resistance development, a selection experiment was performed using the WT, LM and HM strains. We exposed bacteria to increasing concentrations of nitrofurantoin over the course of an eight-day experiment. The increasing selective pressure drives resistance evolution and results in the generation of drug-resistant bacterial isolates. To initiate populations, freezer stocks of WT, LM and HM bacteria were inoculated into 5 ml of MH2 broth (without nitrofurantoin) via an ice scrape. Cultures were left to incubate overnight. After growth, parallel cultures were distributed across six 96-microtitre plates (*n* = 360 populations per strain).

On day 1 of the experiment, MH2 containing 4 μg/ml nitrofurantoin was inoculated with cultures via pin replication from the plate containing the overnight cultures, and subsequently left to incubate overnight. On each subsequent day (days 2-8), fresh nitrofurantoin selective media was inoculated with bacteria via pin replication from the previous days’ plate. During days 1-5, media contained 4 μg/ml (0.25×MIC). The concentration was then doubled daily from day 6 onwards, with bacteria being exposed to 8 μg/ml (0.5×MIC) on day 6, 16 μg/ml (1×MIC) on day 7, and 32 μg/ml (2×MIC) on day 8. OD readings were taken each day to measure bacterial growth and establish population extinction (OD<0.05). At the end of the experiment, surviving populations were stored in MH2 and glycerol (25%) at −80 °C.

### Growth assays of resistant strains

To analyse growth parameters, growth assays were conducted on nitrofurantoin resistant and nitrofurantoin susceptible ancestral WT, LM and HM bacteria. Freezer stocks of isolates were streaked to singular colonies on MH2 agar and left to incubate overnight. The following day, MH2 broth was inoculated with a singular colony from each plate and was left to incubate overnight. Overnight cultures were transferred into a 96-well plate and 1 μl of culture was transferred via pin replication into a plate containing fresh MH2 media. The plate was then immediately placed in the plate reader and left to grow at 37°C for 24 hours, OD readings at 600 nm were taken every 15 minutes.

### Whole genome sequencing of experimentally-evolved resistant isolates

Whole genome sequencing was used to characterize the genetic changes associated with nitrofurantoin resistance in the three genetic backgrounds. A total of 30 experimentally evolved nitrofurantoin resistant isolates (ten isolates from each of the WT, LM and HM backgrounds) were sequenced. Sequencing was performed by MicrobesNG (https://www.microbesng.com, Birmingham, UK) and samples were prepared according to their protocols. Trimmed reads were aligned to the reference genome of *E. coli* str. K-12 substr. BW25113 (Grenier et al. 2014). Mutation calling was performed with breseq (version 0.38.0) (Deatherage and Barrick 2014).

### Genomic analysis of nitrofurantoin-resistant clinical isolates

We analysed the genomes of nitrofurantoin-resistant clinical isolates for variants in DNA replication fidelity and repair genes. A total of 43 isolates were described in two recent papers (Wan et al. 2021; Dulyayangkul et al. 2024). Genomes were obtained from the European Nucleotide Archive (BioProject accessions PRJEB38850 and PRJEB72122). Raw Illumina sequence reads were trimmed using Trim Galore (version 0.6.10, https://github.com/FelixKrueger/TrimGalore), which combines Cutadapt (Martin 2011) and FastQC (version 0.12.0, https://www.bioinformatics.babraham.ac.uk/projects/fastqc/). Trimmed reads were assembled into contigs using Spades (version 4.0.0), (Prjibelski et al. 2020), and assemblies were annotated using Prokka (version 1.14.5) (Seemann 2014).

From the annotated genomes, we extracted amino acid sequences for genes of interest into a single multi-FASTA file per gene. This included nitrofurantoin resistance genes *nfsA* and *nfsB*, and genes implicated in DNA replication fidelity and repair, including the genes involved in LM and HM, *mutS* and *dnaQ*, and others, including *mutH, mutL, mutT, mutY, mutM, nth, ung, uvrA, uvrD* (Fowler and Schaaper 1997; Fowler et al. 2003; Garushyants et al. 2024). To identify variants in these genes, we performed multiple sequence alignment of on translated amino acid sequences using Clustal Omega (Sievers and Higgins 2018). For one isolate, the *mutS* mapped across contigs, which were annotated by Prokka as two separate protein fragments; we additionally used NCBI BLAST on the nucleotide sequence upstream and downstream of the partial *mutS* gene fragments to detect whether its coding sequence had been disrupted, e.g. by an insertion or inversion.

We used two approaches to predict the impact of each variant on protein function. First, we conducted a literature search to determine if specific variants had previously been reported to have a mutator effect. Additionally, we reviewed publications for each gene to assess any known structural or functional importance of the identified residues and the potential for the identified variant to disrupt secondary structure. Second, we used SIFT (Sorting Intolerant from Tolerant) to predict the impact of amino acid substitutions based on sequence conservation (Ng and Henikoff 2003; Sim et al. 2012). SIFT calculates a score for each variant, with scores below 0.05 indicating a predicted deleterious effect on protein function, suggesting potential disruption or loss of activity.

### Calculation of protein disruption scores for *nfsA* and *nfsB* SNVs

For all SNVs identified in *nfsA* and *nfsB* in the experimental and clinical isolates, we also calculated a SIFT score to predict the impact of each substitution on protein function. These scores were used to assess whether the identified SNVs were likely to contribute to nitrofurantoin resistance through loss of function in NfsA or NfsB.

### Statistical analysis

All data handling statistical analyses were performed using the R statistical computing environment (R Core Team 2024) (version 4.4.0) using the tidyverse libraries (Wickham et al. 2019).

## Results

### Increased mutation rate increases the probability of evolving nitrofurantoin resistance in experimental *E. coli* populations

Over the time-course of the experiment, population survival differed markedly between WT, LM and HM populations (**Figure 1A**). A binomial logistic regression indicates differences in survival probability between strains was significant, and that the decline in survival over time was different between WT, LM and HM populations (*χ*^2^ = −96.3, p < 0.0001). WT populations declined sharply over days 6 to 8 (8 μg/ml to 32 μg/ml), with only 28.3% surviving at the final concentration. The largest decrease in WT survival was observed on day 7 of the study, when the MIC of nitrofurantoin (16 μg/ml) was reached. Over the same time-period, a slight decline in survival was observed for LM populations (83.3% surviving), but no decline was observed for HM populations (93.3% surviving).

**Figure 1.**
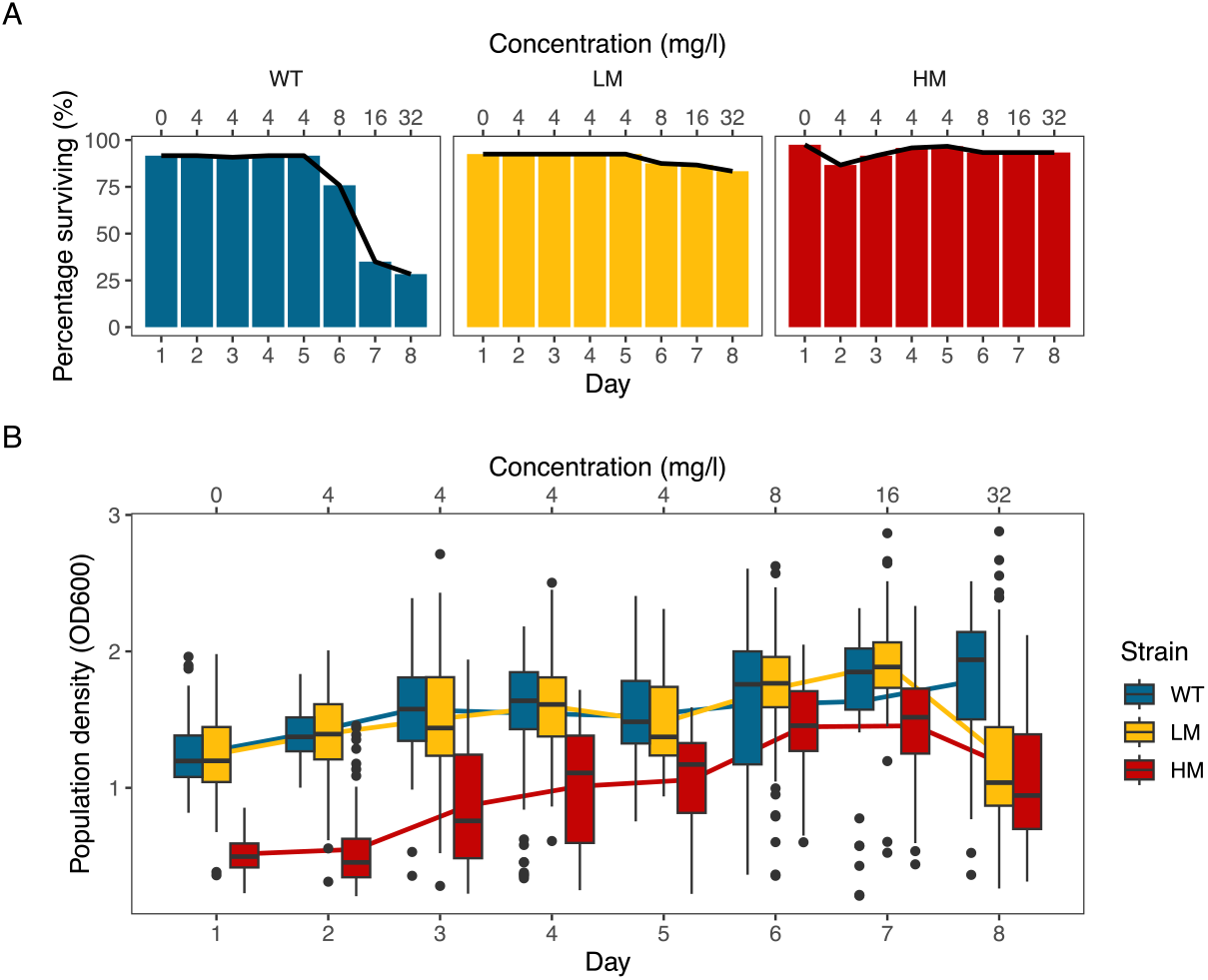
Elevated mutation rates increase the probability of nitrofurantoin resistance evolution, but not the density of surviving populations. **A)** Percentage of surviving populations founded by wildtype (WT), low mutator (LM) and high mutator (HM) strains over the course of the selection experiment, revealing a marked advantage for mutators in surviving nitrofurantoin treatment. **B)** For surviving populations only, surviving WT and LM populations had similar population density across the time-course of the experiment, whereas HM populations had reduced density (OD600—optical density at 600nm).

In contrast, differences in the population density of surviving populations was higher in WT and LM than HM (**Figure 1B**). We analysed this data using linear regression, with strain, day and their interaction as independent variables (Adjusted *R*^2^=0.35, *F*_5,2472_ = 262.5, *p* < 0.00001). The density of surviving populations increased significantly over time (*t*_1_ = 7.7, *p* < 0.00001), despite the increasing antibiotic concentrations. WT and LM populations were consistent in density across the experiment (*t*_2_ = 1.6, *p* = 0.1), and increased slightly less over time than HM populations (*t*_1_ = −2.3, *p* = 0.02). Density of HM populations remained lower than WT populations over the course of the experiment (*t*_2_=17.8 *p* < 0.00001), but did increase more over time than WT populations (*t*_1_=6.3, *p* < 0.0001). At the conclusion of the experiment, although WT populations had lowest probability of survival, the population densities were on par with LM. In contrast, HM, which had greatest probability of survival, had the lowest density across the time-course of the experiment. This was due to an initial growth deficit relative to WT and LM, which was not fully ameliorated this deficit by its conclusion.

### Wild-type and mutator isolates accessed distinct mutational spectra in experimental *E. coli* populations

Whole genome sequencing was performed to identify mutations acquired by experimentally evolved WT, LM and HM isolates (**Figure 2, Table S1**). The three strains exhibited differences in both the number and spectrum of mutations that had been acquired. Across the genome, there were roughly log-fold differences in the number of new mutations appearing in isolates from the WT, LM and HM genetic backgrounds, 3.22 ± 0.972 S.D., 20.7 ± 4.55 S.D., and 376 ± 56.3 S.D. mutations, respectively (**Figure 2**). Genome wide, single-nucleotide variants (SNVs) were more commonly observed in LM and HM isolates compared to WT isolates, which had a more even split between SNVs, mobilisation of insertion sequences (IS), and indels. The bulk of mutations in both LM and HM were transition mutations (i.e. GC>AT and AT>GC), which is typical in strains that have *mutS* deletions. HM had more than twice as many GC>AT as AT>GC mutations (2.46 ± 0.300 S.D.), compared to LM isolates where this ratio was closer to parity (1.14 ± 0.481 S.D.). This change in mutation rate and spectrum reflects the underlying distribution of spontaneous mutations prior to selection, as has previously been characterised for a strain defective in both MMR and DnaQ (Shibai et al. 2023).

**Figure 2:**
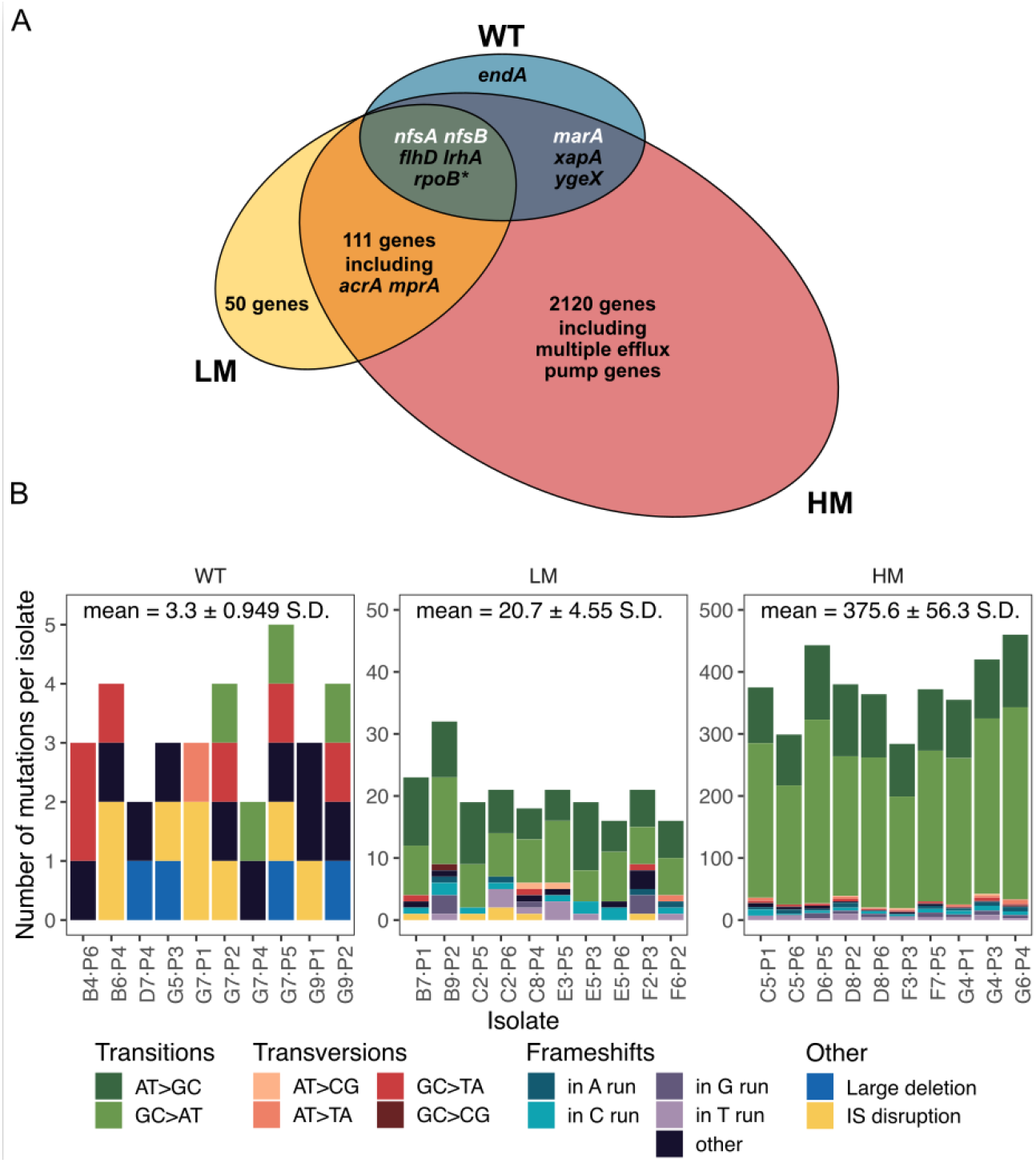
The overlap between genetic targets of nitrofurantoin selection is underpinned by distinct mutational spectra in wild-type and mutator isolates of *E. coli*. **A)** Venn diagram illustrating the overlap of genes mutated in wild-type (WT), low mutator (LM) and high mutator (HM) isolates strains resulting from experimental selection for nitrofurantoin resistance. Genes in white text indicate those specific to nitrofurantoin resistance. **B)** Stacked bar plots displaying the number and types of mutations observed per isolate, showing distinct rates and spectra of mutations in each strain.

Across all three genetic backgrounds, mutations occurred in 2393 genes, more than half of the *E. coli* K-12 genome (**Figure 2B**). Aside from resistance genes, only a handful were found in common between WT, LM and HM isolates, namely *flhD* and *lrhA*, both related to the expression of flagella. These changes likely represent adaptation to the serial transfer protocol (see methods). *xapA* and *ygeX* mutations were also found in both WT and HM isolates, but their functional significance is unknown. LM and HM isolates shared a larger number of genes, which included *acrA* and *mprA* involved in multi-drug resistance. Additional targets common between LM and HM were related to biofilm production, including pilin and fimbriae genes, which may also represent adaptation to the serial transfer protocol. However, the functional significance of the majority of the LM and HM mutations is unclear. Many are likely neutral, as they tended to be synonymous SNVs, conservative non-synonymous SNVs, intergenic (and not in regulatory regions), or in pseudogenes. Their frequency may have increased due to hitch-hiking with resistance mutations.

Additional mutations in multidrug efflux mechanisms were found in LM and HM isolates, but not in WT isolates (**Figure 2B**), indicating that high mutational supply may drive resistance to multiple antibiotics. Mutations occurred in genes encoding major efflux pumps, including EmrAB-TolC, AcrAB-TolC, AcrAD-TolC, and AcrEF-TolC. Loss-of-function mutations in *emrR* (*mprA*), a regulator of *acrAB* and *emrAB*, were also identified in several LM and HM isolates, potentially increasing efflux activity. Previous studies have shown that such mutations can modestly increase nitrofurantoin MIC when combined with *nfsA* and *nfsB* (Puértolas-Balint et al. 2020). Mutations in *tolC* were also observed in HM isolates. The importance of RND and ABC family pumps in this experiment is, however, less clear. Some HM isolates had putatively inactivating mutations in RND family pump genes, including *acrB, acrD* and *acrF*. Non-conservative amino acid substitutions were also seen in ABC genes *macA* and *macB*. MacAB-TolC is reportedly specific for macrolides (Teelucksingh et al. 2020), so it is unclear whether these mutations are functionally significant.

### Mutational spectrum alters the type of resistance mutations occurring in *nfsA* and *nfsB*, but not nitrofurantoin resistance phenotypes

Nitrofurantoin-resistant isolates from all strains had mutations in *nfsA*, and nearly all also had mutations in *nfsB* (**Table 1**). The sole exception was a single WT isolate with a large deletion involving *marA; nfsA* and *nfsB* are both part of the mar-sox-rob regulon, so this likely reduces expression of both genes (Chubiz 2023). There was significant diversity in *nfsA* and *nfsB* mutations, which included SNVs, small indels, large deletions, and IS disruptions (**Figure 3)**. Mutation types varied by strain: WT isolates primarily had IS disruptions and large deletions, while LM and HM isolates showed a higher proportion of SNVs and small indels causing frameshift mutations. This likely reflects differences in their mutational supply between the three strains. For LM, frameshifts in homopolymeric tracts were particularly common, which is characteristic of Δ*mutS* (Foster and Trimarchi 1994; Harfe and Jinks-Robertson 1999). These likely cause inactivation of protein function. For HM, SNVs in residues with characterised roles in protein function or at highly conserved sequence regions. This was particularly evident for *nfsA*, where functionally important residues were mutated in seven isolates. Notable examples were H11Y (CAT→TAT, four isolates), S13F (TCC→TTC, one isolate) and R203H (CGT→CAT, one isolate). H11 and S13F are important for hydrogen bonding between residues, and R203 is involved in the interaction between NfsA and its substrates, i.e. NADPH or nitrofurantoin (Kobori et al. 2001). All of these involve GC>AT mutations, which are elevated in HM.

**Table 1:**
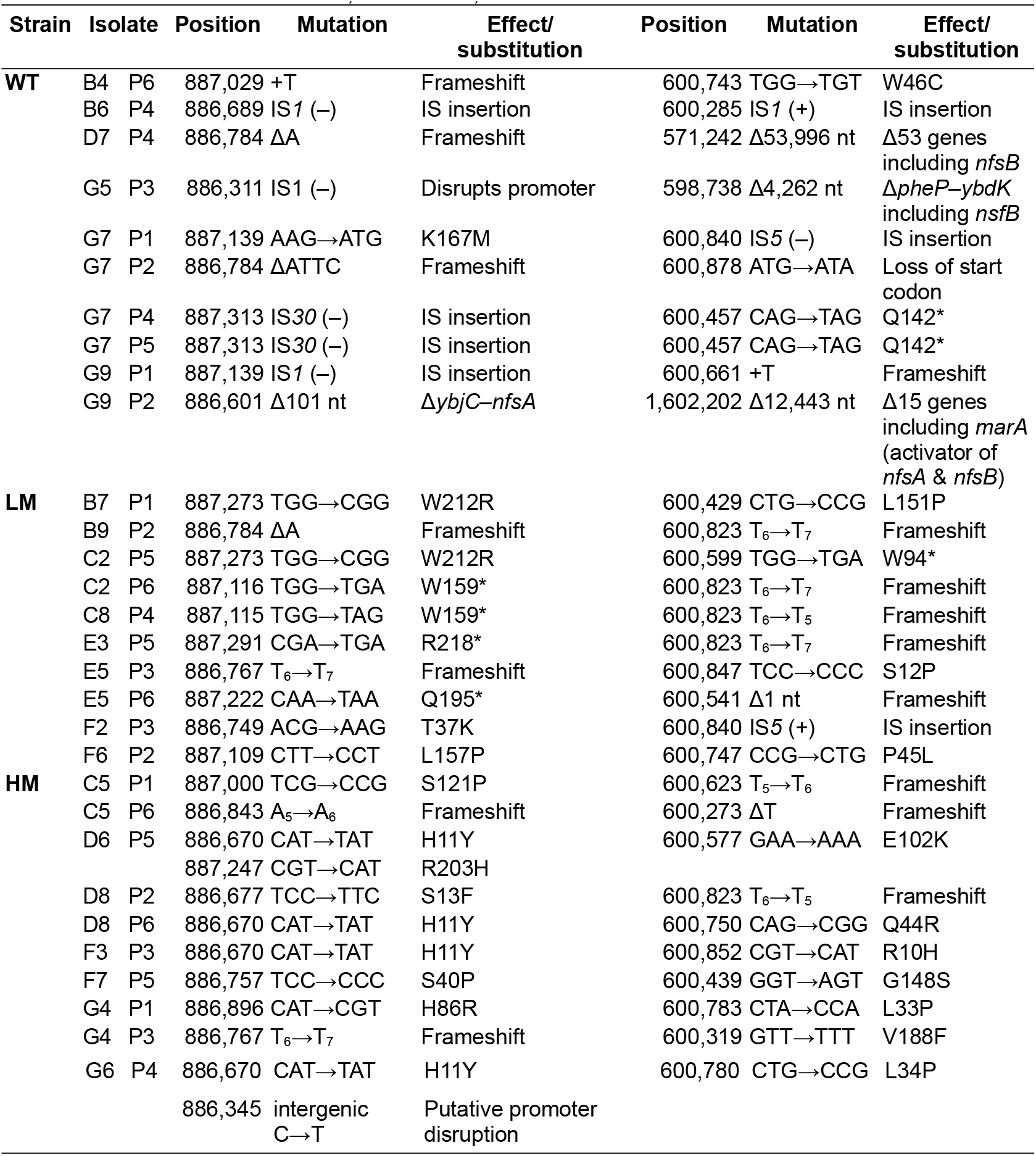
*nfsA* and *nfsB* mutations in nitrofurantoin-resistant lineages of *E. coli* str. K-12 substr. BW25113 experimental evolution. Ten isolates were sequenced from wild-type (WT), low mutator (LM) and high mutator (HM) populations. IS–insertion sequence, (+)/(–)–indicates orientation of IS insertion on forward/reverse strand, Δ—deletion, +—insertion.

**Figure 3:**
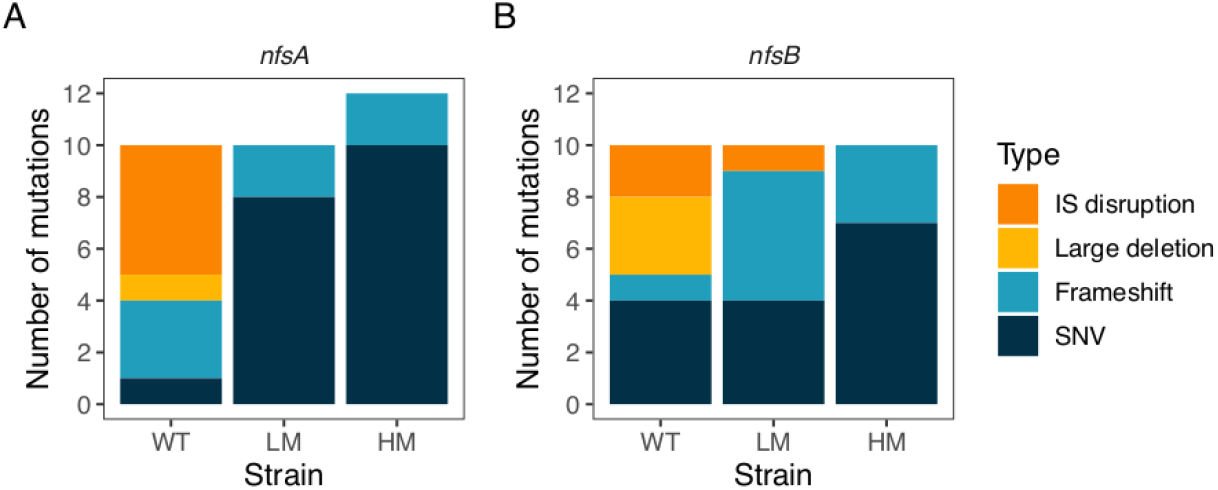
Increased mutation rate alters the types of mutations observed in *nfsA* and *nfsB* in experimentally-evolved nitrofurantoin resistant isolates of *E. coli*. A) Distribution of mutation types in *nfsA*. B) Distribution of mutation types in *nfsB*. IS disruption—disruption of the coding sequence due to an insertion sequence. Large deletion—deletion of 101 nucleotides to 53k nucleotides, potentially involving multiple genes. Frameshift—small insertion/deletion mutations causing a disruption to the reading frame of the gene. SNV—single nucleotide variant.

To determine the level of nitrofurantoin resistance gained by experimentally evolved isolates, MIC assays were performed on both ancestral and experimentally evolved WT, LM and HM isolates (**Figure 4**). Ancestral WT, LM and HM all had the same MIC (16 μg/ml). While lower nitrofurantoin MICs have been reported for strains with defects in different DNA repair systems (nucleotide excision repair and double-strand break repair) (Jenkins and Bennett 1976; Obaseiki-Ebor and Akerele 1986; Ribeiro et al. 2020; Revitt-Mills et al. 2022)), this was not observed for our strains, which have defects in different DNA repair systems. Isolates from the WT populations showed the lowest level of nitrofurantoin resistance (median = 32 μg/ml). Isolates from LM and HM populations gained a higher level of resistance than WT isolates (median = 64 μg/ml). While slightly more HM isolates had a MIC of 64 μg/ml, some LM isolates achieved a higher maximum level of resistance, with a small proportion of LM cultures attaining a MIC value of 128 μg/ml. However, overall, there was no effect of strain on nitrofurantoin MIC values obtained for evolved isolates (one-factor ANOVA, strain: *F*_2,28_ = 0.13, *p* = 0.88). MIC was also not significantly affected by the number of mutations acquired within strains (two-factor ANOVA, strain: *F*_2,21_ = 0.94, *p* = 0.41, mutation count: *F*_1,21_ = 0.05, *p* = 0.83, interaction: *F*_2,21_ = 1.34, *p* = 0.28), nor the presence of mutations in efflux pump systems (two-factor ANOVA, strain: *F*_2,22_ = 0.92, *p* = 0.41, mutation count: *F*_2,22_ = 0.77, *p* = 0.39, interaction: *F*_2,28_ = 0.44, *p* = 0.51).

**Figure 4:**
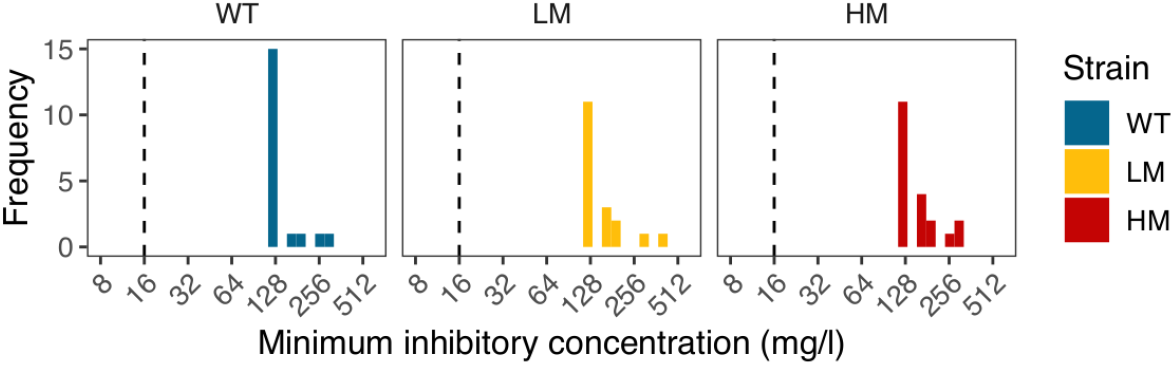
Distribution of minimum inhibitory concentrations (MICs) from experimentally evolved nitrofurantoin-resistant isolates. Resistant isolates arose in populations founded by wildtype (WT), low mutator (LM) or high mutator (HM) strains.

Growth assays were performed on ancestral strains and the sequenced WT, LM and HM resistant isolates (*n* = 10 each), to identify whether mutation rates had an impact on growth of resistant strains in the absence of antibiotic (**Figure 5**). We analysed the effect of strain, and log-transformed mutation count within strain on growth, and found no effect of mutation count within strain (nested ANOVA: strain *F*_2,54_ = 104.9, *p* < 0.0001, log_10_ mutation count within strain *F*_3,54_ = 0.065, *p =* 0.98). Tukey post-hoc comparisons indicated that HM isolates were less fit than both WT isolates (*p* < 0.00001) and LM isolates (*p* < 0.00001), but that WT and LM isolates did not differ (*p* = 0.99). Moreover, WT and LM isolates were indistinguishable from their susceptible ancestral strains, suggesting the absence of a cost of resistance. In contrast, HM isolates exhibited reduced growth relative to WT and LM isolates, but also improved in growth relative to their nitrofurantoin-susceptible ancestral strain. We also estimated growth parameters from the growth curves, i.e., maximum growth rate, carrying capacity, and doubling time (**Figure S1 A-D**). Estimated growth parameters also tended to be greater for WT and LM isolates than for HM.

**Figure 5:**
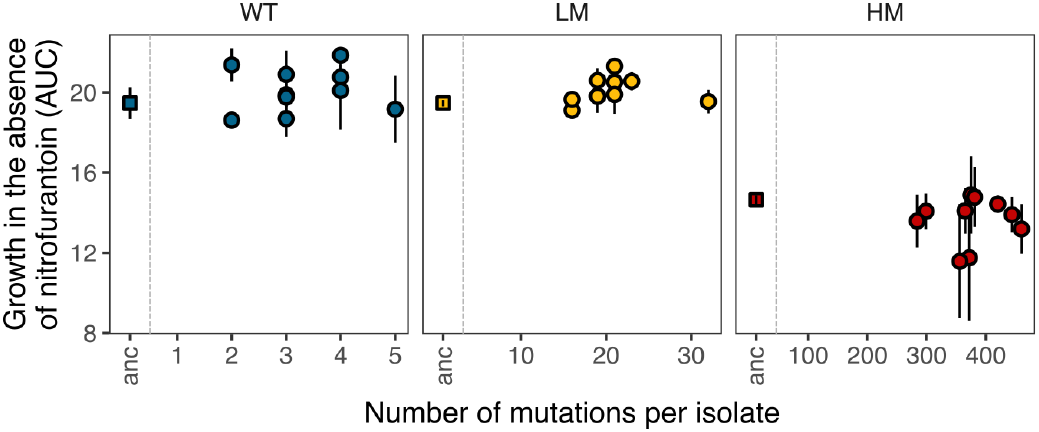
Nitrofurantoin resistance had minimal effects on growth in experimentally evolved isolates (relative to their ancestral strain), irrespective of the number of genomic mutations acquired. Growth was assessed as the area under the curve (AUC) from growth curves measured by optical density at 600 nm (OD600nm). A total of 10 isolates were measured for each strain type: wild-type (WT), low mutator (LM), and high mutator (HM). Additional growth parameters are provided in **Figure S1**.

### Association between nitrofurantoin resistance mutations and variation in DNA replication fidelity and repair in clinical *E. coli* genomes

We analysed the genomes of nitrofurantoin-resistant clinical isolates (Wan et al. 2021; Dulyayangkul et al. 2024) to determine whether resistance co-occurs with variants in DNA replication fidelity and repair genes. Clinical isolates had similar types of mutations in *nfsA* and *nfsB* as observed in our evolution experiment. Notably, IS disruption, while detected in clinical isolates, did not occur as frequently as in the selection experiment. We found variants in DNA replication fidelity and repair genes for 17/43 isolates (39.5%, **Figure 5, Table S2**). These isolates were more likely to have nitrofurantoin resistance encoded by SNVs than those without (*nfsA*: 62.5% vs. 55.6%, *nfsB*: 81.2% vs. 55.6%, **Table S3**), although the patterns were less striking than in our selection experiment and were not statistically significant (Fisher’s exact tests, *nfsA*: *p* = 0.85; *nfsB*: *p* = 0.14). Identified variants involved genes for mismatch repair *(mutS* integrase and IS disruption, *mutH* Q20L and A73G, and *mutL* A350P and A503S), base excision repair (*mutM* P36L), nucleotide sanitation (*mutT* P36S) and DNA polymerase III subunit ε (*dnaQ* A101T). These variants either had a previously-described effect on mutation rates, were computationally predicted to disrupt function by SIFT, or were a substitution known to affect secondary structure in a functionally-important region. The nature of the variants detected is described in **S1 Text**.

### Computational predictions of the impact of *nfsA* and *nfsB* SNVs observed in experimental and clinical isolates

We observed a variety of mutation types in *nfsA* and *nfsB*, some of which clearly disrupt protein function, which is essential for nitrofurantoin resistance. Others, such as SNVs, are less straightforward to interpret and may not contribute to resistance. We therefore used the computational tool SIFT to predict the likelihood that the SNVs would disrupt protein function. SIFT scores for SNVs in *nfsA* (**Table 2**) and *nfsB* (**Table 3**) were analysed for both experimental and clinical isolates. Most experimental isolates had only a single SNV, which was predicted by SIFT to disrupt protein function. However, three SNVs in *nfsB*—L33P, L34P, and E102K—were not predicted to be disruptive. These mutations occurred in α-helices and involve amino acids known to affect helix structure (Javadpour et al. 1999), suggesting they may still impair function. In contrast, many clinical isolates harboured multiple SNVs in both *nfsA* and *nfsB*. Nearly all clinical isolates with SNVs in *nfsA* (11/12) had at least one variant predicted to be disruptive. However, many clinical isolates with multiple SNVs in *nfsB* (11/20) had no individual variant predicted to disrupt function. Additionally, we generated SIFT matrices for all possible amino acid substitutions in both NfsA and NfsB, which can be used to identify potentially disruptive SNVs in other isolates (**Tables S5** and **S6**).

**Table 2:**
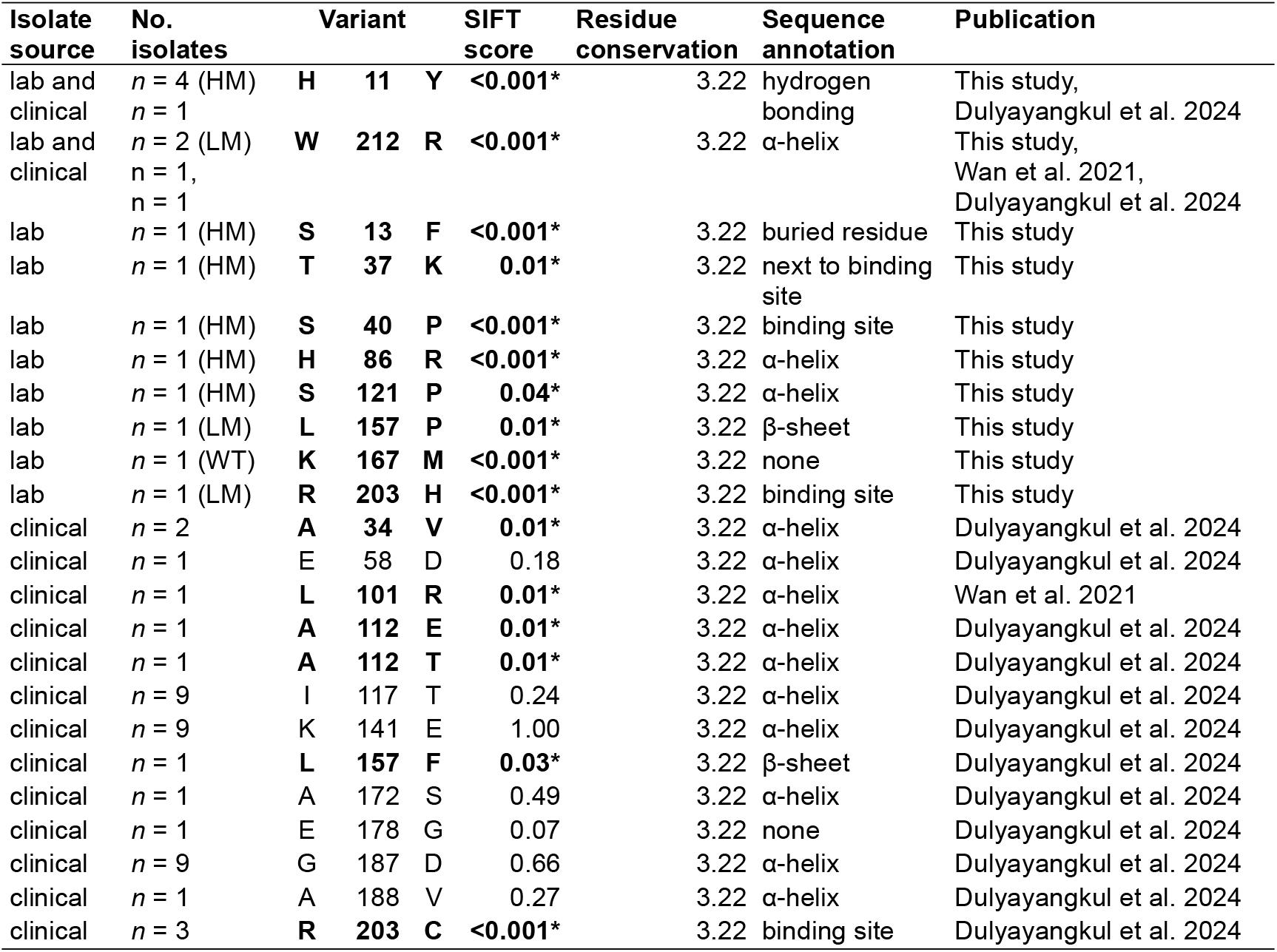
SIFT scores of *nfsA* variants from laboratory experimental evolution and clinical isolates with confirmed nitrofurantoin resistance phenotypes. Residues predicted by SIFT to disrupt function are indicated with * and bold type. Data for individual isolates available in **Table S4**.

**Table 3:**
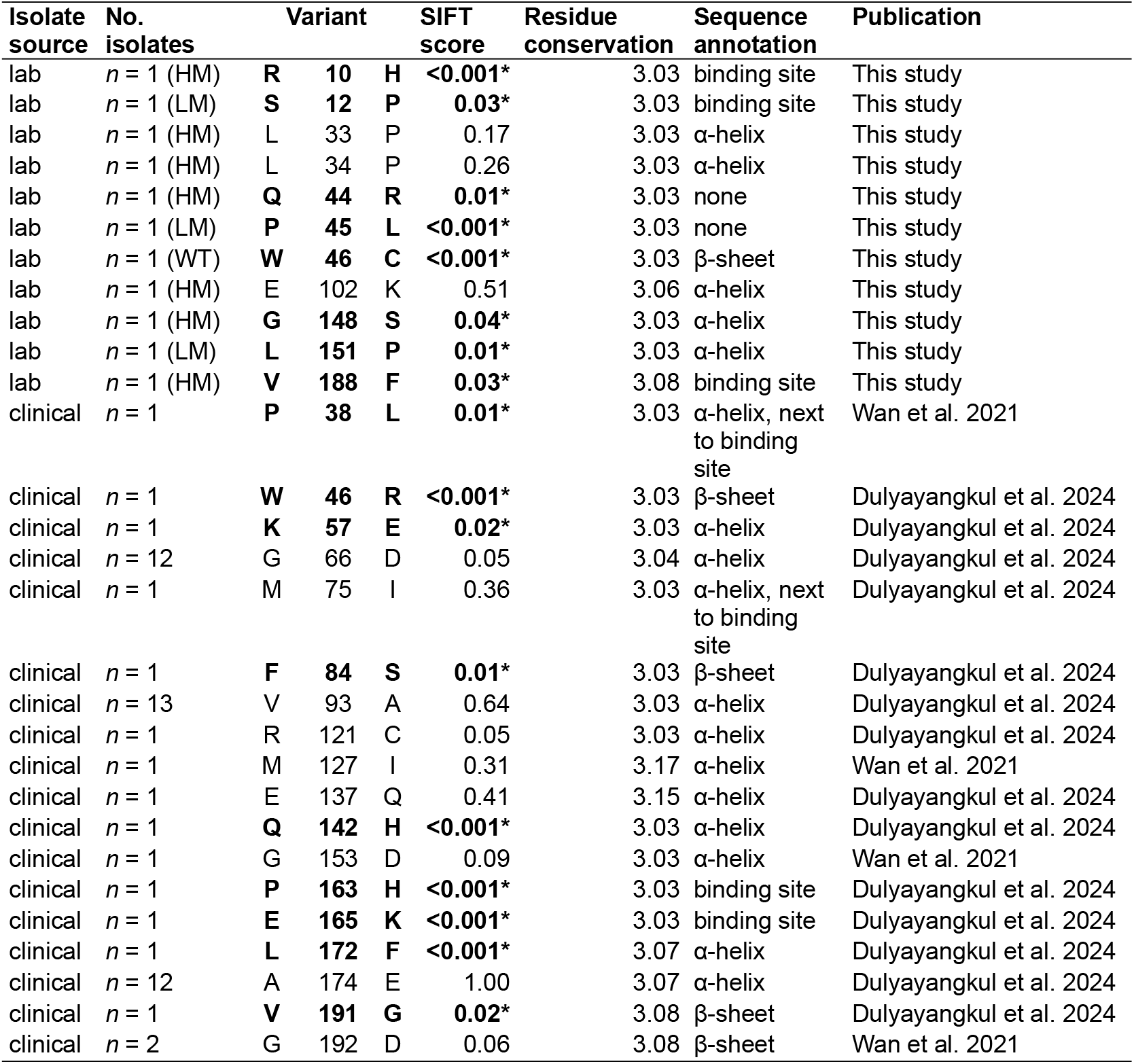
SIFT scores of *nfsB* variants from laboratory experimental evolution and clinical isolates with confirmed nitrofurantoin resistance phenotypes. Residues predicted by SIFT to disrupt function are indicated with * and bold type. Data for individual isolates available in **Table S4**.

## Discussion

Nitrofurantoin has remained remarkably durable against resistance despite sustained use. Resistance to nitrofurantoin develops through spontaneous mutations, but requires two independent mutations in different genes—a challenge for bacteria with wild-type mutation rates, but one that mutator organisms stand a chance of overcoming (Gifford et al. 2023). Using experimental evolution and genomic analyses of clinical isolates, we tested how mutators contribute to nitrofurantoin resistance evolution. Our experiments showed that mutators evolved nitrofurantoin resistance more readily than wild-type strains, with multi-locus mutators particularly prone to resistance emergence (**Figure 1**). Resistance mutations always involved *nfsA* and *nfsB* (**Figure 2**), but their nature varied: wild-type strains acquired broad, disruptive changes (frameshifts, large deletions), while mutators, especially multi-locus ones, gained more precise amino acid substitutions at active sites (**Figure 3, Table 1**). Notably, some mutators also acquired multi-drug resistance, raising broader clinical concerns.

Mutators not only increased the likelihood of nitrofurantoin resistance but also altered the spectrum of resistance mutations. Resistant isolates arising from the WT genetic background tended to have highly disruptive mutations, including small indels producing frameshifts, large deletions and gene disruptions caused by insertion sequences. In the LM background, premature stop codons (often arising from GC>AT transitions) and frameshifts in homopolymeric runs reflected the characteristic mutational signature of Δ*mutS* (Tago et al. 2005). In contrast, HM isolates acquired more ‘refined’ mutations targeting specific active sites in NfsA and NfsB. However, these more targeted changes did not yield higher MICs (**Figure 4**) or improved growth in the presence or absence of nitrofurantoin (**Figure 5**), possibly due to the HM strain’s inherent growth deficit. Whether the mutagenic effects of nitrofurantoin contributed to resistance is unclear. GC>TA transversions, typically elevated by nitrofurantoin, only rarely occurred within *nfsA* and *nfsB*. However, frameshift mutations, which can also be elevated, were relatively common. For LM and HM, it is unclear whether this is solely due to *mutS* deletion or due to synergy between *mutS* deletion and nitrofurantoin.

Among nitrofurantoin-resistant clinical isolates, 39.5% harboured disruptive mutations in DNA repair genes (**Figure 6**), placing them at the higher end of previously reported mutator frequencies for uropathogenic *E. coli* (Denamur et al. 2002; Baquero et al. 2004; Couce et al. 2016). Notably, these mutations included changes in the same genes implicated in our experimental evolution studies, such as *mutS* and *dnaQ*. However, the most frequently observed mutation was a SNP in the nucleotide sanitation gene, *mutT*, which occurred in eight isolates. The phenotypic effects of this variant have yet to be characterised, but it occurs at a highly conserved residue adjacent to the nucleotide recognition motif, and was predicted by SIFT to be disruptive. Mutations in DNA repair genes were associated with an increase in SNVs in *nfsA* and *nfsB*, although the increase was not statistically significant. Future work should consider surveillance for mutators and a more comprehensive phenotypic characterisation of circulating variants in DNA repair genes.

**Figure 6:**
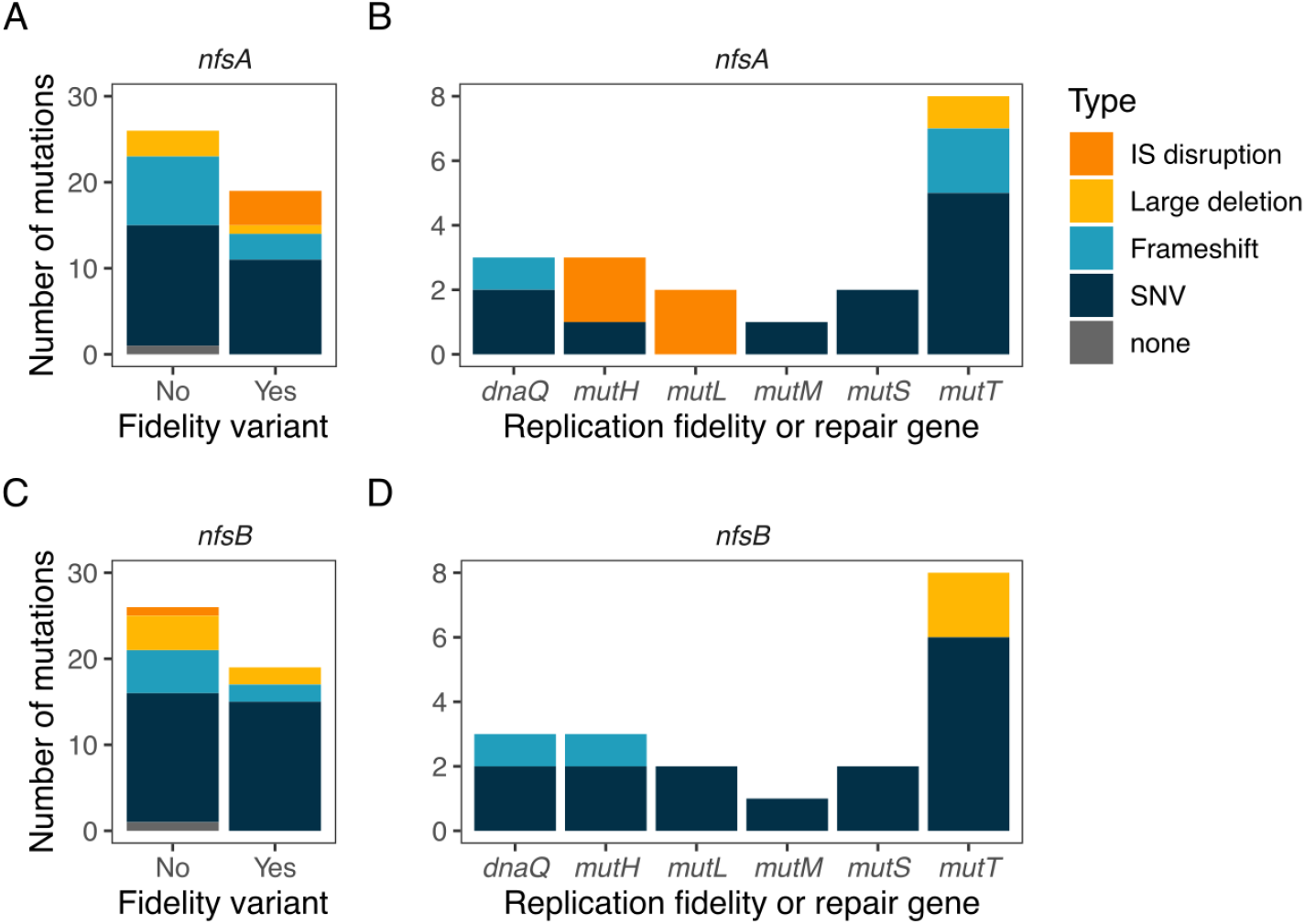
Types of *nfsA* and *nfsB* inactivating mutations nitrofurantoin-resistant clinical isolates with and without variants in DNA replication fidelity and repair genes. **A)** Distribution of mutation types in *nfsA* among isolates with or without variants in DNA replication fidelity and/or repair genes. **B)** *nfsA* mutation types linked to specific DNA replication fidelity and/or repair genes. **C)** Distribution of mutation types in *nfsB* among isolates with or without variants in DNA replication fidelity and/or repair genes. **D)** *nfsB* mutation types linked to specific DNA replication fidelity and/or repair genes. IS disruption—disruption of the coding sequence due to an insertion sequence. Large deletion—deletion of >100 nt, potentially involving multiple genes. Frameshift—small insertion/deletion mutations causing a disruption to the reading frame of the gene. SNV—single nucleotide variant.

The diversity of resistance-conferring mutations in *nfsA* and *nfsB* underscores the difficulty of predicting nitrofurantoin resistance from genomic data (Khamari et al. 2022; Dulyayangkul et al. 2024). Current prediction tools, such as CARD and AMRFinder+, only contain a small fraction of potential resistance mutations. While frameshifts, premature stop codons and insertion sequences usually indicate loss-of-function, predicting the effects of SNVs is more challenging. Several computational tools exist for predicting the impact of amino acid substitutions on protein function (Yazar and Özbek 2021). While none are specifically designed for antimicrobial resistance, they may still offer valuable predictions. Previous studies found PROVEAN scores correlated well with nitrofurantoin resistance (Zhang et al. 2018; Mottaghizadeh et al. 2020; Sorlozano-Puerto et al. 2020; Wan et al. 2021). Here we used SIFT, which predicted disruption for most of the *nfsA* and *nfsB* SNVs in our experimental isolates. However, prediction in clinical isolates with multiple SNVs becomes more difficult when no individual SNV is scored as disruptive. In these cases, disruption may arise from epistatic interactions between multiple SNVs, which cannot be inferred independent-site models like SIFT. Approaches that account for epistasis, such as direct coupling analysis (DCA, (Figliuzzi et al. 2016, Vigué and Tenaillon 2023), could better identify synthetically disruptive SNV combinations.

Beyond nitrofurantoin, mutators present a broader challenge for constraining antibiotic resistance. This is particularly problematic for other antibiotics where resistance is conferred by mutations rather than horizontal gene transfer. Ciprofloxacin is one such antibiotic also used to treat UTIs, where combinations of diverse SNVs in DNA gyrase (*gyrA, gyrB*) and topoisomerase IV (*parC*) are linked to high level resistance, and mutators may have contributed to this (Huseby et al. 2017). Mutators and elevated mutation rates are a problem for managing non-bacterial infections. Mutators have been described for the emerging fungal pathogen *Candida auris*, which has exceedingly high rates of antifungal resistance compared to other *Candida* species (Burrack et al. 2022). High mutation rates associated with RNA viruses have also been implicated in treatment failure (Duffy 2018), and variants have also been discovered in the replicative RNA polymerase of SARS-CoV-2 (Pachetti et al. 2020). Addressing the role of mutators, and high mutation rates more generally, is therefore crucial for developing strategies to curb the spread of treatment resistant infectious disease.

## Conclusion

Collectively, our results indicate that mutators could affect nitrofurantoin resistance in multiple ways. Elevated mutation rates not only increase the likelihood of resistance emerging but also facilitate the development of more specific resistance mechanisms tailored to nitrofurantoin without broadly impacting other genes. Given the wider evidence linking mutators to antibiotic resistance, future research should prioritise examining the role of mutators in clinical infections and their connection to the emergence of nitrofurantoin resistance. This is particularly important as mutation-driven resistance may develop during the course of treatment. The switch to nitrofurantoin as a first-line choice against UTIs, in the UK and elsewhere, may exacerbate this problem. Future strategies to preserve the efficacy of nitrofurantoin should focus on monitoring the prevalence of mutators in UTIs, along with enhancing surveillance to detect emerging resistance.

## Acknowledgements

This work was supported by the Academy of Medical Sciences (Springboard Grant SBF007\100096), and UKRI-BBSRC (BB/X007979/1). The authors would like to thank Mato Lagator for comments on a draft of the manuscript. RK would like to thank CD Durrant for assistance with R programming.

## Data availability statement

All data are available from Figshare (doi:10.48420/27119436) and were made available during the review process (https://figshare.com/s/b2d9c35f8e20037c9790). Genome sequences generated in this study are archived at the European Nucleotide Archive (ENA, BioProject accession PRJEB80644). Genome sequences of clinical nitrofurantoin-resistant isolates were made available by the original authors at the ENA (BioProject accessions PRJEB38850 and PRJEB72122).

## Author contributions

Conceptualisation: DRG. Initial manuscript draft: RK, DRG. Performed experiments: RK, JF. Analysed data: RK, DRG. Supervisory roles: DRG, JF. Initial manuscript draft: RK. Edited and approved the manuscript: all authors.

## Figures

**Figure S1:**
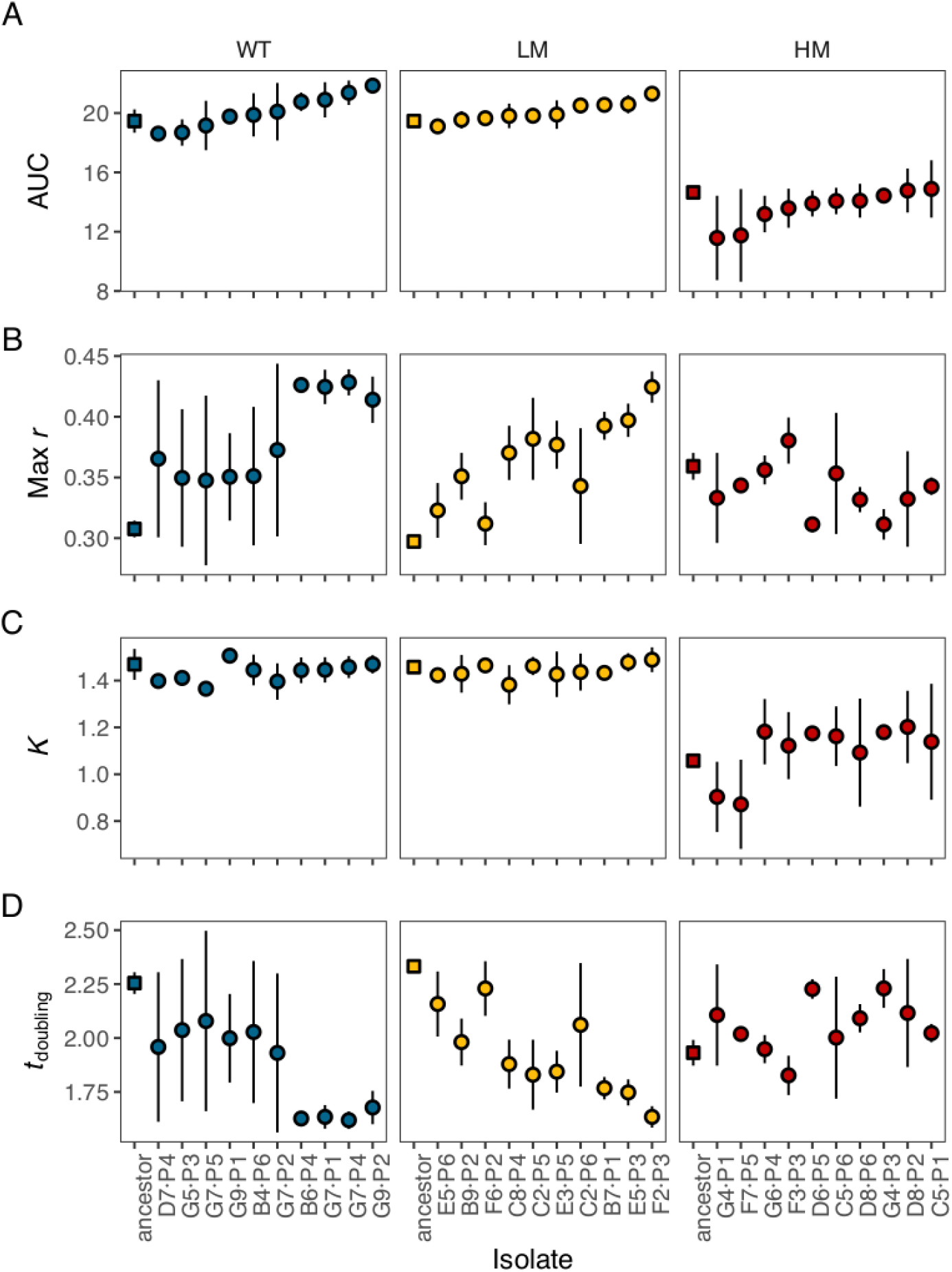
Growth parameters of experimentally evolved nitrofurantoin-resistant isolates from genetic backgrounds with different mutation rates. Parameters were estimated by fitting a logistic curve to growth curves measuring optical density at 600 nm (OD600nm). A) AUC—area under the curve. B) Max *r*—maximum growth rate. C) *K*—carrying capacity. D) *t*_doubling_—doubling time. Resistant isolates arose in populations founded by wildtype (WT), low mutator (LM) or high mutator (HM) strains. Isolates are ordered on the horizontal axis according to increasing values of AUC in panel A.

