## Supplementary Text for "Mutators can increase both the rate of occurrence and specificity of nitrofurantoin resistance mutations"

### Evolutionary risk analysis of mutators for the development of nitrofurantoin resistance

#### Descriptions of variants found in DNA replication fidelity and repair genes

We analysed the genomes of 43 nitrofurantoin-resistant clinical isolates (Wan et al. 2021; Dulyayangkul et al. 2024) to assess whether resistance is associated with variants in DNA replication fidelity and repair genes. We performed multiple sequence alignment using Clustal Omega (Sievers and Higgins 2018) on the protein sequences of several genes implicated in DNA replication fidelity and repair, including and, and others, including *mutS* *mutH*, *mutL*, *mutY*, *mutM*, *mutT*, *dnaQ*, *nth*, *ung*, *uvrA*, *uvrD* (Fowler and Schaaper 1997; Garushyants et al. 2024). Putatively disruptive variants were found in *mutS* *mutH*, *mutL*, *mutM*, *mutT*, and *dnaQ*. A discussion of their functional significance is provided below.

##### Mismatch repair *mutS*, *mutH*, *mutL*

The *mutS*, *mutH*, and *mutL* genes together form the MutSHL holoenzyme, a crucial component of the DNA mismatch repair (MMR) pathway that identifies and corrects replication errors such as mismatched bases and insertion-deletion loops. Mutations in any of these genes can disrupt the holoenzyme's function, compromising the cell's ability to maintain genomic stability. This impairment can lead to an accumulation of mutations across the genome, contributing to a mutator phenotype that significantly increases the likelihood of developing antibiotic resistance. We found distinct variants in 11 isolates in genes encoding the mismatch repair holoenzyme, MutSHL.

Two variants were detected in the mismatch recognition component, *mutS.* In one case, *mutS* was disrupted by an IS*3*-like insertion sequence (*n* = 1 isolate). A second variant involved an unusual N-terminal sequence with an integrase gene in the vicinity of the promoter (*n* = 1 isolate), neither of which occur in its closest nitrofurantoin-sensitive relative, strain EC958 (Forde et al. 2014).

Two variants were observed in the strand recognition component *mutH*, Q20L (*n* = 2 isolates) and A73G (*n* = 1 isolate). Q20L is predicted to disrupt function by SIFT. A73G was not predicted by SIFT to disrupt function, but occurs in an *α*-helix within the D(X)6–30(E/D)XK endonuclease motif (Ban and Yang 1998). Glycine is known as a '*α*-helix-breaker' (O'Neil and DeGrado 1990), and so may disrupt function.

In *mutL*, two variants were detected at position 350, but were not predicted to disrupt function by SIFT. The consensus residue is alanine, but some isolates possessed a proline or serine instead (A350P, *n* = 1 isolate; A350S, *n* = 5 isolates). We note that *E. coli* str. K-12 possesses an S at position 350. This variant therefore is unlikely to have a strong mutator phenotype, though we note that in a study of mutation rates in diverse *E. coli*, *E. coli* str. K-12 substr. MG1655 had a higher than average mutation rate (Richards 2019).

##### Base excision repair *mutM mutY*

A single variant was found in the base excision repair gene *mutM* (*n* = 1 isolate). The *mutM* gene encodes a formamidopyrimidine-DNA glycosylase, which plays a crucial role in the base excision repair pathway by removing oxidatively damaged guanine residues (8-oxoguanine) from DNA, thereby preventing mutagenesis. A P36L substitution in *mutM* could have significant functional consequences, as indicated by the SIFT score. Proline usually has a role in protein folding and stability, and substitution with leucine could disrupt MutM conformation or its ability to interact with DNA. However, the specific variant has not been previously described in the literature.

##### Nucleotide sanitation *mutT*

A putatively disruptive variant was found in *mutT,* P36S (*n* = 7 isolates). MutT is responsible for scavenging highly mutagenic oxidized GTP molecules (8-oxo-dGTP). Knockouts of *mutT* have a high mutation rate to antibiotic resistance (Krašovec et al. 2017; Emam et al. 2024). While P36S has not yet been functionally characterised, residue 36 is immediately adjacent to the nudix motif, which is required for 8-oxo-dGTP recognition (residues 37-59). Altering residue 37 increases mutation rate, though not to the same extent as complete deletion of the gene (Emam et al. 2024). The proline at position 36 is highly conserved across bacteria and archaea, which is also suggestive of its functional importance.

**DNA polymerase III subunit ε *dnaQ***

A single variant in the same gene as our HM strain was detected, *dnaQ* A101T (*n* = 3 isolates). A101T has been previously linked to a mutator phenotype (Luan et al. 2013), although SIFT did not predict that A101T would be disruptive. Similar to T15A in HM, A101T occurs within an exonuclease region crucial for proofreading, specifically ExoII (Whatley and Kreuzer 2015).
